# Characterization Standard for *In-situ* Cryo-electron Tomography

**DOI:** 10.64898/2026.05.20.726049

**Authors:** Mallak Ali, Joshua Hutchings, Tahiti Dutta, Nikki Jean, Garret Greenan, Elizabeth A. Montabana, Jonathan Schwartz, M.G. Finn, Matthias Haury, David A. Agard, Bridget Carragher, Mykhailo Kopylov, Mohammadreza Paraan

## Abstract

Standardized biological specimens are essential for optimizing cryoEM workflows and benchmarking instrument performance. While apoferritin fulfills this role for single-particle analysis, no equivalent exists for cryo-electron tomography. Ribosomes are frequently used but require large datasets due to C1 symmetry and structural heterogeneity, limiting rapid optimization and standardized comparison of workflows. Here, we present PP7 virus-like particles (VLPs) overexpressed in *E. coli* as a scalable in situ benchmark. VLPs have high orders of symmetry enabling rapid, high-resolution validation of tomographic pipelines from minimal datasets, while their distinct structural features across low to high resolutions provide a practical resolution metric.

## Main

Cryo-electron tomography (cryoET) provides three-dimensional visualization of macromolecular assemblies within their native cellular environment^1^. In combination with subtomogram averaging (STA), cryoET can reach subnanometer and, in some cases, near-atomic resolution^2^. Despite rapid advances in specimen preparation, data acquisition, processing, and reconstruction algorithms, the field lacks a widely adopted benchmark specimen for evaluating resolution of in situ tomography workflows^3^. Ribosomes have frequently been used for testing cryoET workflows because of their high abundance and contrast^4,5^. However, the lack of symmetry and heterogeneity of ribosomes means that a decent resolution (∼4 Å) ribosome structure using STA generally requires a large dataset (many tens^2^ to hundreds^4,5^ of tomograms). This presents a bottleneck in cases where many conditions need to be rapidly compared, or when microscopy experiments are challenging, such as when testing novel hardware approaches. A highly symmetric structure reduces dataset size requirements, one of the reasons that apoferritin (Octahedral) has been a good benchmark for single particle analysis (SPA). However, the featureless shell of apoferritin precludes the estimation of intermediate resolutions thus limiting in situ benchmarking. Here, we demonstrate that PP7 bacteriophage virus-like particles (VLPs), readily overexpressed at high yield in *E. coli*, provide a robust benchmark that can be used to rapidly test in situ cryoET workflows (Figure 1). Their large size (2.5MDa, ∼26 nm diameter), high symmetry (Icosahedral), and distinct structural features across all resolution ranges (Figure 1c), meet all the requirements of an efficient cryoET benchmark. They can be prepared both by plunge freezing, or as focused-ion beam milled (FIB-milled) lamellae after high-pressure freezing, both methods suitable for small or large-scale cryoET data collection (Figure 1a).

**Figure 1.**
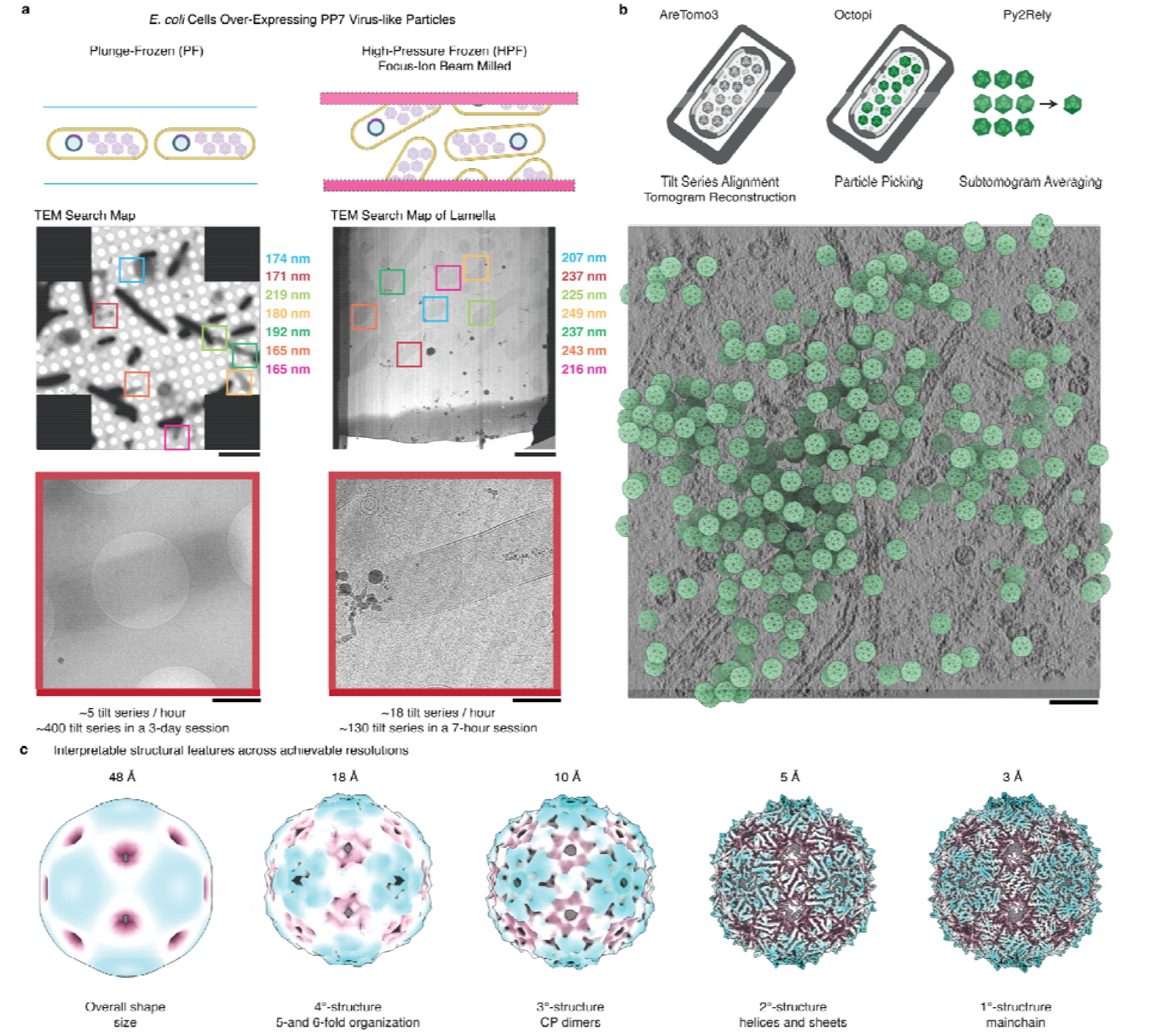
Expression of PP7 VLPs in *E. coli* provides a reproducible benchmark specimen for in situ cryoET workflows. **a**. Two methods of vitrification for PP7 induced *E. coli*: plunge freezing (PF) or high-pressure freezing (HPF). In PF, a monolayer of cells decorates the grid (TEM Search Map, top) with many thin electron-transparent cells (TEM Search Map, bottom). For HPF, a 20-micron-thick bulk of cells is milled using the Waffle^12^ method (TEM Search Map of Lamella). Color-coded squares are indicated with corresponding tomogram thickness values. **b**. General scheme of data processing steps. The tomograms reveal a high abundance of intracellular PP7 assemblies (green overlaid maps). **c**. Isosurface renderings of the PP7 VLP reconstructions (colored based on the distance from the center) demonstrate robust signal across multiple spatial frequencies. Left to right: At 48 Å, the overall size and shape are resolved. At 18 Å, the global icosahedral morphology can be inferred. At 10 Å, individual Coat Protein (CP) dimers and their organization into pentamers and hexamers becomes apparent. At 5 Å, secondary structural elements, including alpha-helices and beta-hairpins, are resolved. At 3 Å, the protein backbone and bulky side-chains can be traced. Scale bars: TEM Search Map-top 10 μm, TEM Search Map-bottom 1 μm, TEM Search Map of Lamella-top 2 μm, TEM Search Map of Lamella-bottom 500 nm, Tomogram 50 nm.

For the plunge-frozen samples, 290 tilt series provided a 3.0 Å (Nyquist) structure using an STA pipeline based on AreTomo3^6^/Octopi/Relion5^7^ without classification (Figure 2a). We further show that we can use only two tomograms to extract more than 2,000 particles and yield a 3.8 Å structure (Figure 2b). At this resolution, the beta-sheets of the capsid protein can be resolved verifying the reported resolution. For the FIB-milled lamellae we processed 124 tilt series using the same pipeline, to achieve a map at 3.9 Å resolution (Figure 2a). Processing a pair of tomograms from FIB-milled lamella, closely matching the plunge-frozen pair in terms of thickness, CTF resolution, and CTF score (Supplementary Table 1), resulted in a 4.2 Å structure (Figure 2b). The lower resolution for the lamellae dataset may be attributed to issues such as charging^8^ or the FIB-induced damage layer^4,9,10^ and will be the subject of larger, more systematic studies. Importantly, because this analysis can be performed using as few as two tomograms, the benchmark enables rapid and quantitative comparison of multiple specimen preparation, data acquisition, and processing workflows across experimental conditions and microscopy facilities.

**Figure 2.**
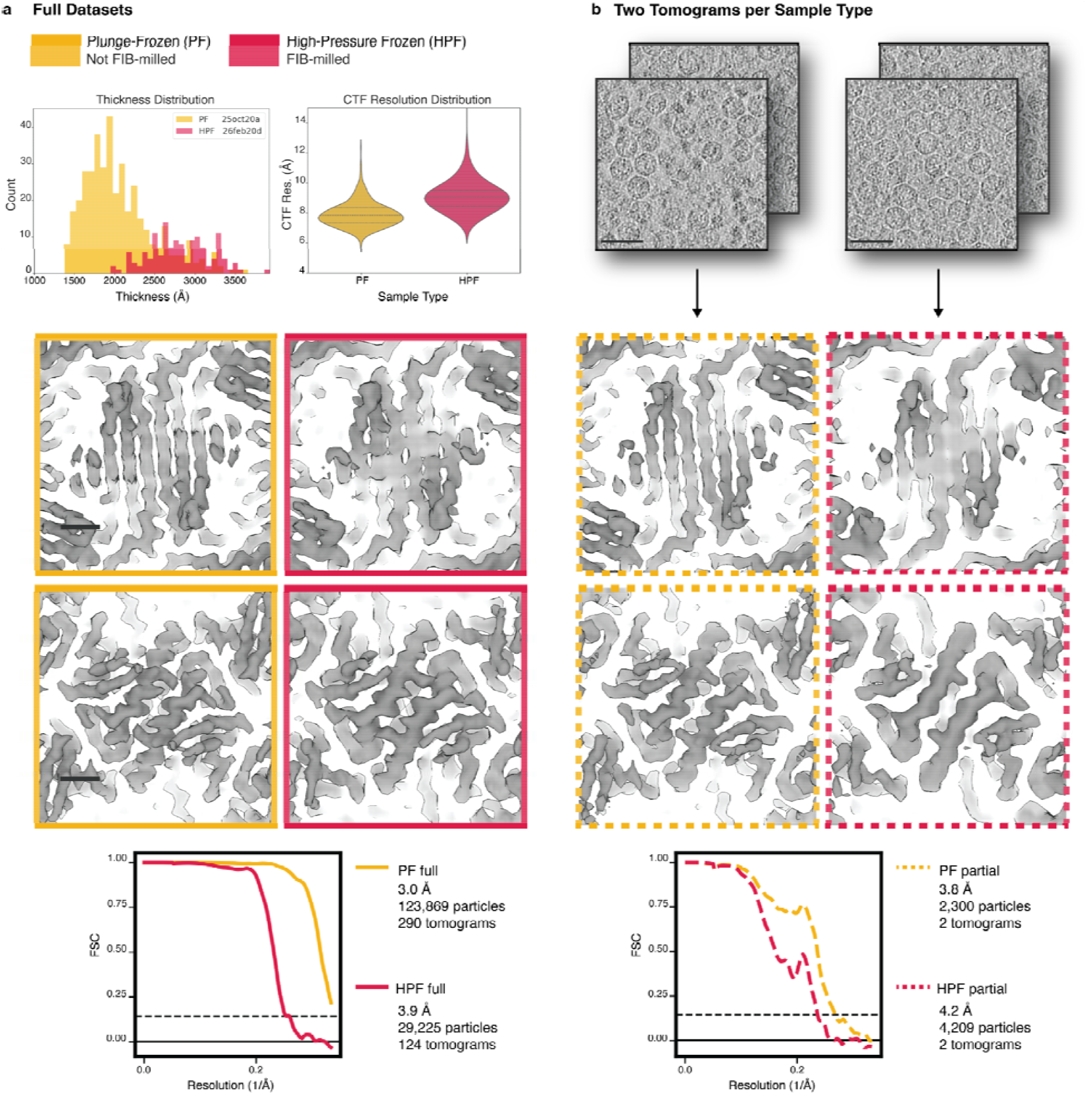
VLPs in *E. coli* benchmark is a resolution standard with plunge-frozen datasets acting as a control for lamellae datasets. **a.** Histogram plots of tomogram thickness distribution as estimated in AreTomo3 for the plunge-frozen dataset and the high-pressure frozen dataset indicate ranges of 150-350 nm and 200-350 nm, respectively. The CTF resolution distributions are: PF: mean=7.9 Å, std=0.8 Å; HPF: mean=9.2 Å, std=1.6 Å. T=3 VLPs from 290 tomograms for the PF dataset and 124 tomograms for the HPF dataset were processed using subtomogram averaging in Relion 5^7^. The PF dataset reaches a Nyquist resolution of 3.0 Å and the HPF dataset reaches a resolution of 3.9 Å. The differences in resolution are highlighted by comparing Beta-sheets (top row) and alpha helices (bottom row). **b**. For a controlled comparison of the PF and HPF datasets, two tomograms from each dataset with matching quality metrics and thicknesses (Supplementary Table 1) were processed using STA de novo. This small PF dataset reaches a resolution of 3.8 Å while the HPF dataset reaches 4.2 Å. The difference is notable in the separation of the beta-sheets and the backbone of the alpha-helices. Maps are displayed as isosurfaces at 5 standard deviations from the mean map value. Scale bars: panel **a** 10 Å, panel **b** 50 nm.

We also collected an untilted dataset of a plunge-frozen sample to test the results of a standard SPA pipeline. By upsampling the EER movies, we overcame the Nyquist resolution limitation (3 Å) achieving 2.89 Å resolution (Extended Data Figure 1). A dose titration experiment shows that with a dose lower than 5 e^-^/Å^2^ per tilt it is not possible to get a high resolution structure of the VLPs (Extended Data Figure 2). Given that the tomography datasets were collected at 3.87 e^-^ /Å^2^ per tilt, we conclude that tilts other than the zero-degree angle are contributing signal to the high-resolution STA structures. Interestingly, we found that we were only able to classify the dominant T=3 VLPs in the SPA datasets. Other geometries^11^ were not resolved in 2D classification (Extended Data Figure 3) by SPA but this was not a limitation in the STA processing. Perhaps this indicates a potential method to illustrate the power of tomography in resolving complex heterogeneity.

Overall, we demonstrate that PP7 VLPs expressed in *E. coli* provide a practical resolution benchmark for in situ cryoET workflows. Unlike previous benchmarks using ribosomes, VLPs enable rapid, quantitative assessment of performance across the whole workflow, without requiring large datasets. By lowering the barrier to systematic optimization and enabling controlled comparisons across experimental variables, this benchmark has the potential to accelerate method development and improve reproducibility across the field.

## Acknowledgements

We thank Shawn Zheng and Ariana Peck at Biohub for their close collaboration in developing the preprocessing pipeline. We are grateful to Yue Yu, David Dong, Anchi Cheng, Michael Souza, and Pallavi Khedle for developing processing and project management tools, and Dari Kimanius for consulting on subtomogram averaging and analysis. We thank Utz Ermel and the CryoET Data Portal team for uploading the datasets. We thank the HPC team at Biohub for support on computation and Gorica Margulis and Armando Soria for lab support and management. This work is funded by Biohub; T.D. and M.F. were supported by the NIH (U54 AI 170855)

## Contributions

M.P., M.K., and B.C. developed the idea for the benchmark sample. T.D. developed the plasmid in M.G.F.’s lab and provided *E. coli* samples for preliminary experiments. M.A. developed the sample preparation and the freezing. N.J. carried out the FIB-milling. G.G. oversaw the FIB-SEM operations. E.A.M. and M.P. collected the tomography and single tilt datasets. M.K. processed the single particle datasets. J.S. and J.H. developed the subtomogram averaging pipeline. M.P. and J.H. processed the tomography datasets and carried out the analysis. All authors contributed to writing and making the figures and tables. M.H., D.A.A. and B.C. provide overall leadership for projects at Biohub, Redwood Shores.

## Data availability

The tilt series datasets have been deposited on the CZ CryoET Data Portal (https://cryoetdataportal.czscience.com/) under Deposition ID: CZCDP-10336. The plunge-frozen dataset and the high-pressure frozen lamellae dataset are DS-10493 and DS-10494, respectively. All maps are publicly available through the Electron Microscopy Data Bank (EMDB) with accession codes: 77085 (plunge frozen full dataset), 77086 (high pressure frozen full dataset), 77087 (plunge frozen partial dataset), 77088 (high pressure frozen partial dataset) and 77092 (plunge frozen by single particle analysis).

## Methods

### Sample preparation

#### PP7 VLP expression and purification

PP7 plasmids^11^ were acquired from the Finn group at Georgia Institute of Technology. BL21(DE3) competent *E. coli* cells were thawed and 50 µL were gently transferred to a 1.5 mL Eppendorf tube. Then, 2 µL of 50 ng/μL PP7 plasmid was added, mixed, and placed on ice for 10 minutes. The mixture was heat shocked at 42°C for 30 seconds, recovered on ice for 5 minutes and transferred to a 14 mL round-bottom tube containing 950 µL of SOC media. After 60 minutes of incubation at 37°C with shaking at 200 rpm, 100 µL of the cell culture was spread onto a streptomycin-selection agar plate and incubated overnight at 37°C. A colony from the selection plate was picked up and used to inoculate 50 mL of 2YT starter media supplemented with 50 µg/ml of streptomycin. It was incubated at 37°C shaking at 200 rpm for 16 hours. Two large cultures of 500 mL of 2YT media were prepared, each supplemented with 50 µg/ml of streptomycin and 10–20 mL from the overnight starter culture. Cultures were incubated with shaking at 200 rpm at 37°C, and optical density at 600 nm (OD_600_) values were checked every 30 minutes. Once an OD_600_ of 0.6 was reached, protein production was induced with 100 µL of 1M IPTG. In one hour intervals up to 4 hours after IPTG addition, 20 mL of culture was collected in duplicate from each large flask, placed at 4°C for 10 minutes, and centrifuged at 3075g (Sorvall X Pro Series Centrifuge TX-1000) for 15 minutes. The supernatant was discarded and the pellets were stored at -80°C for long-term storage. The datasets described in this paper were collected on cells at 1 hour and 2 hours post induction for the plunge-frozen 25oct20a and FIB-milled 26feb20d, respectively (Extended Data Figure 5).

### Grid Preparation

#### Vitrification

Samples were vitrified using two different methods: plunge freezing and high-pressure freezing. Pellets from the time-point of interest were thawed by hand until no ice was visible.

#### Plunge freezing

The pellet was resuspended in ∼200 µL of LB broth and transferred to a 1.5 mL Eppendorf tube. OD_600_ was measured and adjusted until a final optical density of 0.6 was achieved. All steps were done on ice. An OD of 0.6 was found to be optimal for creating a thin monolayer of bacteria on the grid, permitting direct imaging in TEM without FIB-milling. Quantifoil R 2/1 Au 200 mesh grids were used, glow-discharged at 15 mA for 45 seconds on the carbon side (Ted Pella Pelco easiGlow). The Leica GP2 plunge-freezer was prepared with the following conditions: 90% humidity, 4°C chamber temperature, 30-second delay before blotting, and 4-second blot time. Then, 4 µL of sample was applied to each grid. The grids were plunged into liquid ethane after blotting. Grids with vitrified cells were clipped with standard TFS Autogrid rings before loading into Krios for data acquisition. Dataset 25oct20a was collected on plunge-frozen grids.

High-pressure freezing: a Leica EM ICE high-pressure freezer was used to establish the waffle protocol^12,13^. To achieve a high concentration of cells, the pellet was gently resuspended in 150 µL of LB broth, to get an OD of ∼100 (i.e. a very viscous pellet). The suspension was transferred to a 1.5 mL Eppendorf tube and centrifuged at 1,500 g for 15 minutes to reform the pellet. To prepare for high-pressure freezing, for every grid, two Leica Type B planchettes with a diameter of 6 mm were sanded using P800, P7000, and P15000 grit in increasing order and cleaned with metal polish. The planchettes were then coated with a solution of 0.5% cetyl palmitate in diethyl ether and placed on Whatman filter paper to dry; a thin film was visible on the planchettes after evaporation. Quantifoil R 1/4 SiO_2_ Au 200 mesh grids were glow-discharged at 15 mA for 45 seconds on the SiO_2_ side. Immediately before high-pressure freezing, the supernatant of the centrifuged sample was discarded leaving a pellet volume of ∼100 μL. Then 10 μL of 100% glycerol was added to the pellet. Then, 3 µL of the pellet was applied per grid. After freezing, grids were gently removed from the planchette hat by pushing tangentially until they slipped off, then clipped with notched TFS Autogrids with the sample side facing down in the clipping station. Dataset 26feb20d was collected on the waffle grids.

### FIB-Milling

#### Cryo-FIB milling of VLPs in e-coli

High-pressure frozen VLPs in *E. coli* were loaded into a Helios Hydra plasma cryo-FIB/SEM microscope and were milled using the Waffle^12,13^ method (Supplementary Table 2). The grid was coated with a layer of Pt-GIS for two minutes, using xenon gas, then sputter coated for two minutes with an ion beam voltage of 12.00kV. The grid was then rotated perpendicular to the FIB to create two trenches in the middle of grid squares that were free of surface debris. Trenches measure 20.00 μm in x, 35.00 μm in y (equivalent to the final milling direction) and were milled with a beam current of 60 nA using Xenon plasma, leaving a slab in the middle that is milled further to produce the final lamella. After trench milling, the grids were then coated for two minutes and thirty seconds with a secondary layer of GIS to add additional protection to the surface during the preparation steps of cleaning and notch milling. A new tile set was created in ThermoFisher software Maps to indicate the location of the trenches.

In the ThermoFisher software AutoTEM (v. 2.4.3), the trenches were linked to lamellae locations and the first step in the preparation tab, eucentric alignment, was completed. Following the alignment, the slab was milled from the bottom to a thickness of 3.5 μm. Then “notch” patterns were manually milled to create stress relief for the lamellae. After creating the notches, the lamellae went through the automated pipeline in AutoTEM with a final thickness set to 120 nm. To produce thinner and more uniform lamellae, the final polishing step (Polishing Step 2 in AutoTEM) was rerun multiple times with manual checks.

### TEM data acquisition

All datasets were collected on a Krios G4 at 300 keV, equipped with a Falcon 4i direct electron detector, a SelectrisX energy filter, and an X-FEG. The datasets were collected using TFS Tomo5 software. All datasets were collected at a calibrated pixel size of 1.50 Å/px (nominal pixel size of 1.54 Å/pix as reported in the mdoc metadata files), nominal defocus of 3 µm and a total dose of 120 e^-^/Å^2^ that was equally spread over 31 tilts spanning -45° to +45° with 3° increments. The energy filter slit width was set to 10 eV. The dose-symmetric strategy was used for collecting the tilt-series. For plunge-frozen cells, tilt images were acquired in .eer format and 234 .eer frames were grouped in bins of size 10. For lamella, tilt images were acquired in .eer format and 225 .eer frames were grouped in bins of size 15. The single particle dataset was collected at a total dose of 60 e^-^/Å^2^ in .eer format, converted to 57 frames on micrograph import and upsampled to a pixel size of 0.77 Å/px, with a nominal defocus range of 2-3 µm. 402 tilt series were collected for the plunge-frozen grid. 135 tilt series were collected for the lamellae. 434 untitled micrographs were collected for SPA analysis.

## Data processing

### Movie and tilt series alignment

The frames were motion-corrected and the tile series were aligned using AreTomo3^6^ (Supplementary Table 3). For tilt series alignment, AlignZ is not specified because the thickness of the sample is estimated automatically by AreTomo3^6^. For the comparison of thickness and CTF values between the PF and HPF datasets, both datasets were processed with the latest version (2.3.0) as noted in Supplementary Table 3.

All datasets were denoised using denoisET^6^ version 0.1.0 and using a model (https://github.com/apeck12/denoiset) that was trained on a synaptosome dataset (https://cryoetdataportal.czscience.com/depositions/10313).

### Particle Picking

For picking VLPs in all datasets, one model was trained in Octopi16 (https://chanzuckerberg.github.io/octopi/). For this training, 3 tomograms from the dataset 25aug25a (https://cryoetdataportal.czscience.com/datasets/10455) were manually picked for the spherical VLPs resulting in 1,521 manual picks. Octopi was then used in model exploration mode to find the best Unet architecture for picking VLPs. The best model was then used to pick all the particles for all datasets. The model is available at 10.5281/zenodo.20057531. The picks were exported to copick^14^ (https://copick.github.io/copick/) for the subsequent steps of STA.

### Sub-tomogram averaging

All STA was carried out using Py2Rely (https://github.com/chanzuckerberg/py2rely), a python workflow for Relion5^7^ built on top the CCPEM pipeliner framework (https://ccpem-pipeliner.readthedocs.io/en/latest/).

#### Sub-tomogram averaging pipeline for dataset 25oct20a

AreTomo3^6^ (v2.2.7) was used for motion correction, tilt series alignment, and CTF estimation. ∼131,000 octopi picks from 290 tomograms were imported into py2rely without any 2D or 3D classification. An automatic pipeline was set up by py2rely. The first refinement was reference-based and was done at a binning of 6 in a Relion^7^ Class3D job type. The reference was a down-sampled icosahedral T=3 map from the phantom dataset^15^ (https://cryoetdataportal.czscience.com/depositions/10310). The resulting map from this first refinement job was used as the reference map for all the subsequent jobs. Because of the dataset’s high quality, no 2D or 3D classification was needed, and a T=3 VLP was refined to 3.02 Å with 3 rounds of polishing at high resolution (bin 1, 1.5 Å/pix).

#### Sub-tomogram averaging pipeline for dataset 26feb20d

AreTomo3^6^ (v2.3.0) was used for motion correction, tilt series alignment, and CTF estimation. Only for this dataset, the tilt series alignment used sample recentering (available only in v2.3.0) and sample tilt correction. This improved the tomogram reconstruction quality which was confirmed visually. It also increased the total number of Octopi picks and the population of the T=3 icosahedral VLPs (Extended Data Figure 4). Octopi picks (∼57800 total from 124 tomograms) were 2D-classified using the slab method^15^, and the classes corresponding to T=3 icosahedral VLPs (∼35,700 picks) were selected for downstream 3D refinement. An automatic pipeline was set up by py2rely. The extraction of subtomograms in Relion was limited to the first 60 electrons (half of the full dose) to avoid the high tilt angle artifacts in the dataset. The first refinement was reference-based and was done at a binning of 6 in a Relion Class3D job type. The reference was the down-sampled icosahedral T=3 map from the phantom dataset. The resulting map from this first refinement job was used as the reference map for all the subsequent jobs. After the first round of full resolution refinement (bin 1, 1.5 Å/pix), CTF refinement, and Bayesian polishing, T=3 VLPs reached 5.20 Å resolution (half-map FSC). These particles were then classified without alignment into two classes, with one class containing 12% of the particles distinct by the presence of RNA bound inside and the other class with 88% of the particles without RNA. The second class of particles were further refined and polished 5 rounds to a resolution of 4.15 Å. Afterwards, VLPs that fell within 30 nm from the top and the bottom of the lamellae were filtered out due to being in the damage layer. This was done by evaluating the coordinates of the center of the VLPs against the thickness measurements of AreTomo3. It is worth noting that all the tomograms were reconstructed to 240 nm, even though the thickness range is 200-350 nm. So for any tomogram thicker than 270 nm, the damage layer was effectively already filtered out from the beginning of the pipeline (particle picking). Therefore, only 1,700 particles were removed by applying the 30-nm threshold. This improved the resolution to 4.03 Å with two rounds of polishing and one refinement in-between. Extending the damage layer to 45 nm did not improve the resolution any further.

#### Sub-tomogram averaging pipeline for two-tomogram subsets

Two tomograms were manually selected from both plunge-frozen and milled high-pressure-frozen datasets based on approximately similar characteristics, such as tomogram thickness, CTF resolution and score (Supplementary Table 1). An automated py2rely pipeline was applied in the same way as outlined above for the full plunge-frozen dataset. An additional five rounds of CTF refinement, Bayesian polishing and 3D refinement were applied to maximize attainable resolution.

#### FSC calculation

The same soft-edge mask obtained from the VLP asymmetric subunit was used for resolution comparison between all maps resolved by STA, using *relion_postprocess*^*7*^ and phase-randomization at 20 Å. Resolution improvements as a function of high-resolution refinement cycles for all datasets are shown in Extended Data Figure 6.

### Single Particle Analysis

#### EER upsampling (Extended Data Figure 1)

The SPA processing was done fully in CryoSPARC v4.1^16^. The EER movies were upsampled to a pixel size of 0.77 Å/px. Movies were corrected with Patch Motion Correction and CTF was estimated with PatchCTF. Initial particles were picked from the first 50 micrographs using the blob picker with the diameter settings of “min 250 Å” and “max 350 Å” resulting in ∼12,000 picks. Particles were extracted with a box size of 640 px and downsampled to a box size of 480 px with a binned pixel size of 1.027 Å. The particle stack was processed in 2D classification and a single well-defined 2D class was selected to be used as a template. Template picking on the full dataset of 434 micrographs yielded ∼120,000 particle picks that were extracted with the box size of 640 pixels downsampled to 480 pixels. After the first round of 2D classification, the particles were filtered to ∼50,000 picks and after the second round further filtered to a final stack of 34,368 particles. An initial model was generated using an ab initio refinement with default settings. Non-uniform refinement was done with icosahedral symmetry and included iterative per-particle CTF defocus refinement and global CTF parameters refinement produced a final map with reported GSFSC_0.143_ resolution of 2.89 Å and estimated Guinier B-factor of 95.

#### Electron titration experiments (Extended Data Figure 2)

The EER frames were grouped such that each frame group had a dose of 1 e^-^/Å^2^. The dose titration experiment was then done by using 4, 5, and 10 frames corresponding to accumulated doses of 4, 5, and 10 e^-^/Å^2^, respectively. We determined that a dose of 5 e^-^/Å^2^ is the minimum dose necessary for the refinements to converge.

## Supplementary Tables

**Supplementary Table 1.** AreTomo3 metrics for the two-tomogram tests for the plunge-frozen and high-pressure frozen datasets. Tomograms were selected to have broadly similar metrics such as estimated thickness, CTF fit resolution and score, and number of bad patches from AreTomo3 local alignments. Alpha0 and Beta0 represent the tilt offset about the tilt axis and orthogonal axis, respectively.

| Sample Type | Plunge-frozen |  | FIB-milled |  |
| --- | --- | --- | --- | --- |
| Tilt Series | Position_142.mrc | Position_58.mrc | Position_9_11.mrc | Position_9_2.mrc |
| Thickness (nm) | 2190 | 2160 | 2190 | 2160 |
| Tilt Axis | -96.06 | -96.50 | -96.52 | -96.03 |
| Global Shift (pix) | 76.50 | 246.94 | 140.11 | 95.69 |
| Bad Patch All | 0.00 | 0.00 | 0.00 | 0.00 |
| Bad Patch Low | 0.00 | 0.00 | 0.00 | 0.00 |
| Defocus (Å) | 27704 | 27638 | 25093 | 24052 |
| CTF-Res (Å) | 7.92 | 7.92 | 8.00 | 7.68 |
| CTF-Score | 0.21 | 0.22 | 0.19 | 0.19 |
| Alpha0 | -2.20 | 4.20 | 22.80 | 22.80 |
| Beta0 | -11.20 | 3.00 | -3.00 | -12.30 |
| Particles | 1218 | 1082 | 2206 | 2003 |

**Supplementary Table 2.** Table of beam currents, patterns, and run times used for each milling step. Waffle milling steps and corresponding beam currents, pattern type, and milling time & depths are listed. Patterns for trench, undercut, and notch milling were manually drawn with the special pattern dimensions listed. Undercut milling was completed by drawing and milling rectangles at three decreasing angle intervals (40, 30, and 20 degrees) to remove the ice until ∼3.5um remained. Patterns for the remaining milling steps were presets in the AutoTEM software. Additionally, ‘polishing 2’ was run two supplemental times after the automated steps were completed to ensure the lamella had a smooth and consistently thin appearance.

| Milling Step | Beam Current (Xe) | Pattern Type | Time/Depth |
| --- | --- | --- | --- |
| Trench Milling | 60 nA | Rectangle<sup>1</sup> | ~30-90 s per trench |
| Undercut Milling | 4.0 nA | Rectangle | ~15 s per milling angle (40,30,20) |
| Notch Milling | 1.0 nA | Rectangle<sup>2</sup> | 120 s |
| Rough Milling | 1.0 nA | Rectangle | 200% Depth Correction |
| Medium Milling | 1.0 nA | Rectangle | 200% Depth Correction |
| Fine Milling | 0.1 nA | Rectangle | 150% Depth Correction |
| Polishing 1 | 30 pA | Cleaning Cross Section | 100% Depth Correction |
| Polishing 2 | 10 pA | Cleaning Cross Section | 70% Depth Correction |

<sup>1</sup> Special Pattern Dimensions - Two rectangles placed 30.00 $\mu\text{m}$ apart: x = 20.00 $\mu\text{m}$ , y = 35.00 $\mu\text{m}$ .
<sup>2</sup> Special Pattern Dimensions - Five rectangles: vertical rectangles - x = 0.25 $\mu\text{m}$ , y = 10.00 $\mu\text{m}$ , horizontal rectangles - x = 0.25 $\mu\text{m}$ , y = 3.50 $\mu\text{m}$

**Supplementary Table 3.** List of AreTomo3 parameters for motion correction and tilt series alignments for the different datasets. PF 25oct20a was used as input for particle picking and subtomogram averaging. PF 25oct20a (run2) was only used for thickness and CTF values distribution in comparison with HPF 26feb20d. HPF 26feb20d was used for particle picking and STA.

| Parameter | PF 25oct20a | PF 25oct20a (run2) | HPF 26feb20d |
| --- | --- | --- | --- |
| Align | 1 | 1 | 1 |
| AlignZ | 0 | 0 | 0 |
| AmpContrast | 0.07 | 0.07 | 0.07 |
| AtPatch | [4, 4] | [4, 4] | [3, 3] |
| CorrCTF | [1, 15] | [1, 15] | [1, 15] |
| Cs | 2.7 | 2.7 | 2.7 |
| DarkTol | 0.7 | 0.7 | 0.7 |
| EerSampling | 2 | 2 | 2 |
| FlipGain | 1 | 1 | 1 |
| FlipVol | 1 | 1 | 1 |
| FmInt | 10 | 15 | 15 |
| Group | [2, 4] | [2, 4] | [1, 3] |
| McBin | 2.0 | 2.0 | 2.0 |
| McIter | 15 | 15 | 15 |
| McPatch | [4, 4] | [4, 4] | [4, 4] |
| McTol | 0.1 | 0.1 | 0.1 |
| Outlmod | 1 | 1 | 1 |
| PixSize | 1.5 | 1.5 | 1.5 |
| ReconRange | [-90.0, 90.0] | [-90.0, 90.0] | [-90.0, 90.0] |
| SplitSum | 1 | 1 | 1 |
| TiltAxis | [0.0, 1.0] | [0.0, 1.0] | [0.0, 1.0] |
| TiltCor | [0.0, 0.0] | [1.0, 0.0] | [1.0, 0.0] |
| VolZ | 1600 | 1600 | 1600 |
| Version | v2.7.0 | v2.3.0 | v2.3.0 |

## Extended Data Figures

**Extended Data Figure 1.**
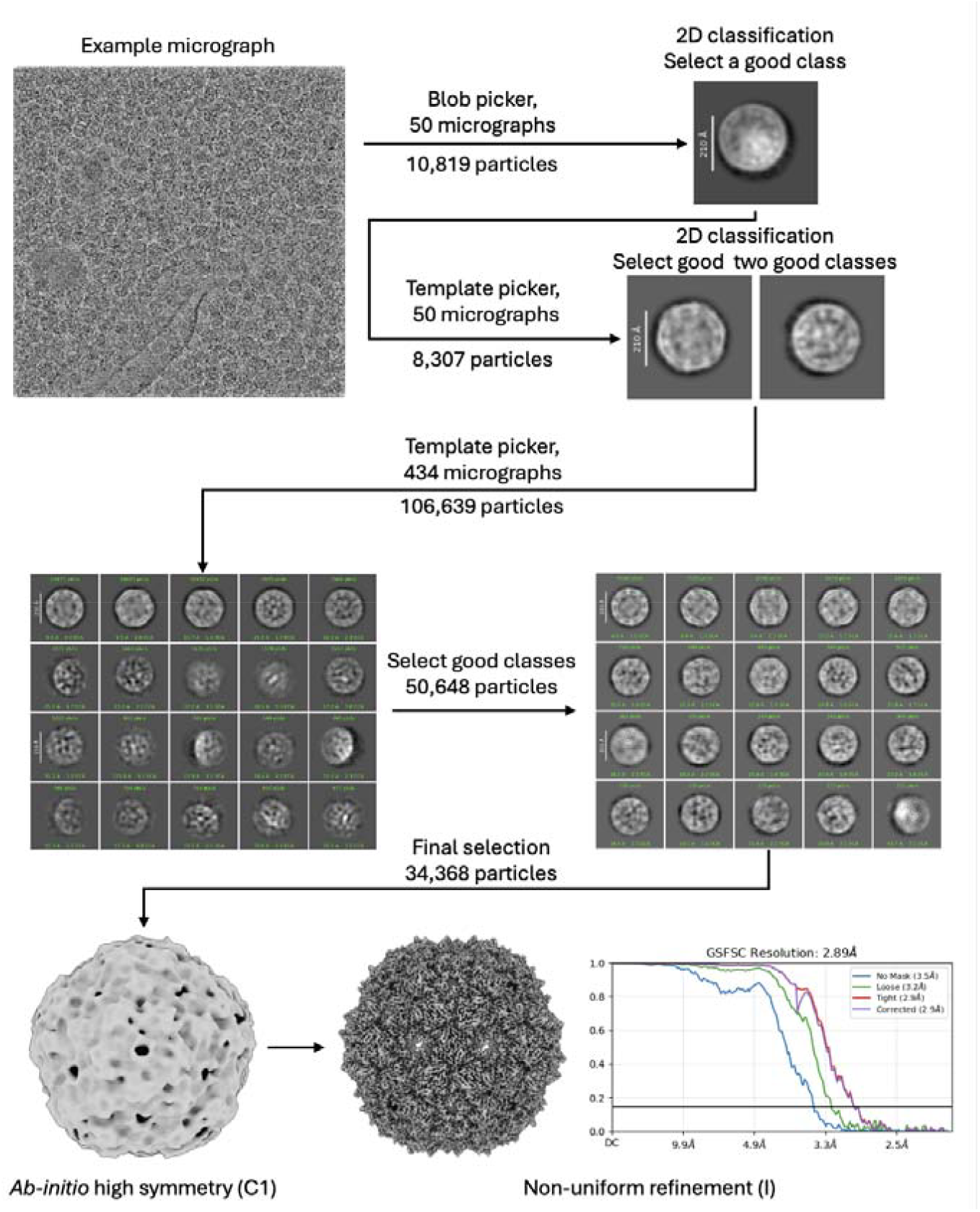
Single-particle analysis (SPA) workflow and high-resolution reconstruction of PP7 VLPs. Representative cryo-micrographs of plunge-frozen *E. coli* cells overexpressing PP7 were subjected to a hierarchical particle selection strategy. An initial template was generated from a subset of 50 micrographs via blob picking and 2D classification to generate a template for template-based picking across the full dataset. Two successive rounds of 2D classification were used to curate a final stack of 34,368 particles. An ab initio model generated without symmetry served as the initial volume for non-uniform refinement. By imposing icosahedral symmetry and employing per-particle CTF and global aberration corrections, the final reconstruction reached a resolution of 2.89 Å. The resulting map shows clear secondary structural definition and bulky side chain densities, consistent with the resolution.

**Extended Data Figure 2.**
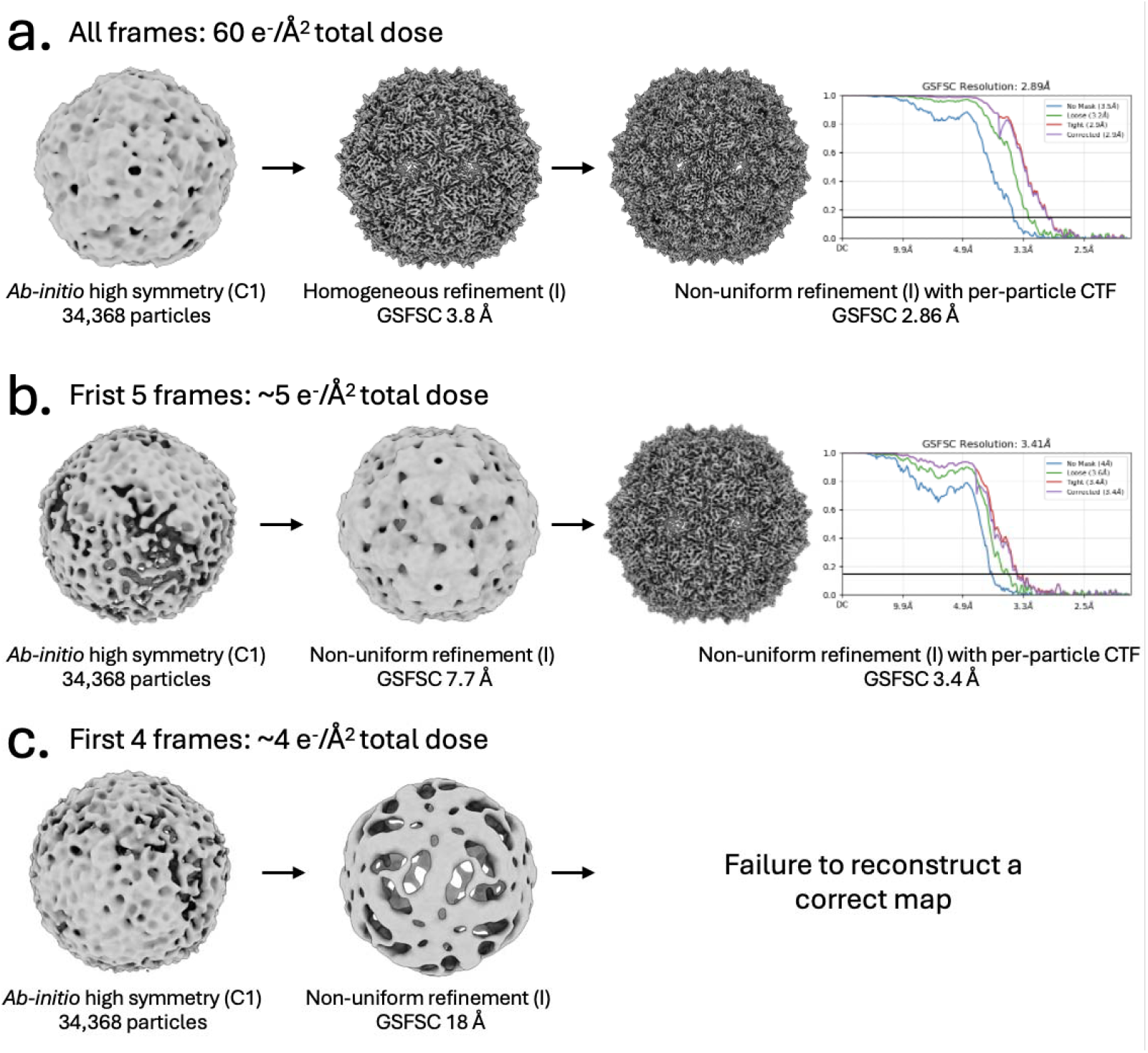
Comparison of dose-limited and particle-limited processing of the plunge-frozen dataset. **a**. Full dataset processed using all movie frames - 34,368 particles. An *ab-initio* model generated without imposed symmetry (C1) was refined using homogeneous refinement followed by non-uniform refinement with per-particle CTF correction, yielding a final reconstruction at 2.86 Å resolution. **b**. Frame-limited dataset processed using first five frames to simulate dose-limited (total dose ∼5 e^-^/Å^2^) conditions. An *ab-initio* model (C1) was used in non-uniform refinement, followed by non-uniform refinement with per-particle CTF correction, resulting in a final reconstruction at 3.4 Å resolution. **c**. Frame-limited dataset processed using first four frames to simulate dose-limited (total dose ∼4 e^-^/Å^2^) conditions. An *ab-initio* model (C1) was used in non-uniform refinement, but failed to produce a correct map.

**Extended Data Figure 3.**
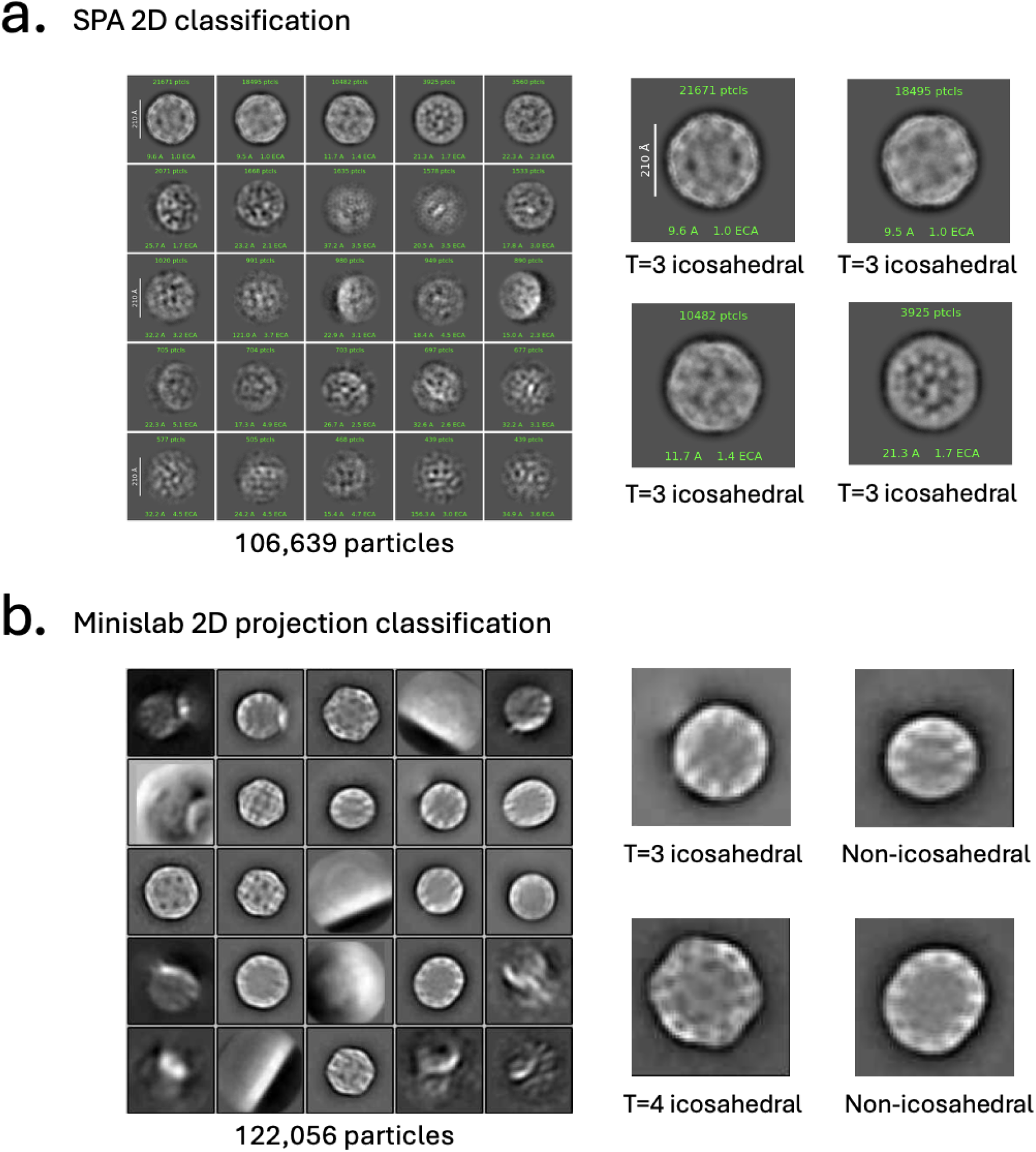
Different geometries of VLPs are resolved in 2D classes of projected subtomograms but not resolved in 2D classes from SPA data. **a**. The single particle dataset was 2D classified in CryoSPARC with T=3 icosahedral VLPs resolved. Other geometries were not resolved despite multiple classification strategies. **b**. The VLP picks from tomograms are projected to 2D images (minislabs^15^) and then 2D classified using Relion 5. Multiple geometries are resolved in this case.

**Extended Data Figure 4.**
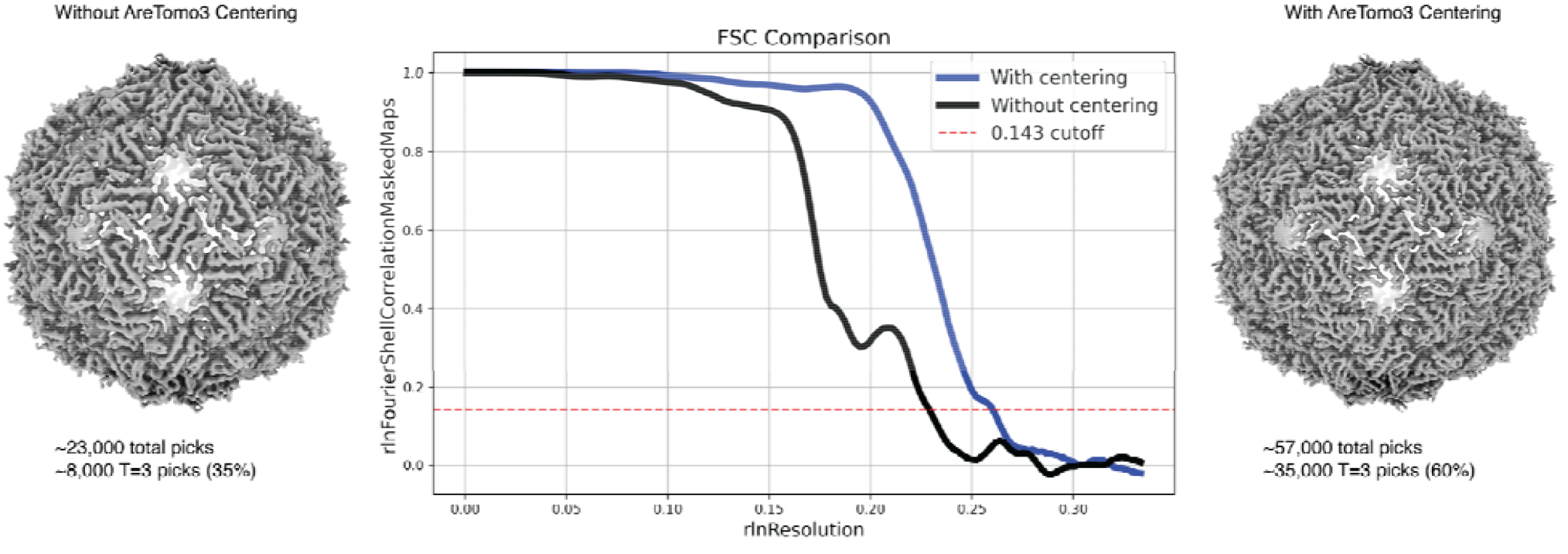
VLPs in *E. coli* benchmarking of AreTomo3 tilt series alignment. On the left, AreTomo3 (v.2.2.7) alignments were done without recentering the sample during alignment. The result was ∼8,000 T=3 icosahedral VLPs reaching a resolution of 4.3 Å. On the right, AreTomo3 (v2.3.0) alignments were done on the same dataset with recentering the sample during alignment. Besides visual improvements in the tomogram reconstructions, more particles were picked (∼35,000 T=3 VLPs) and higher resolution of 3.9 Å was achieved.

**Extended Data Figure 5.**
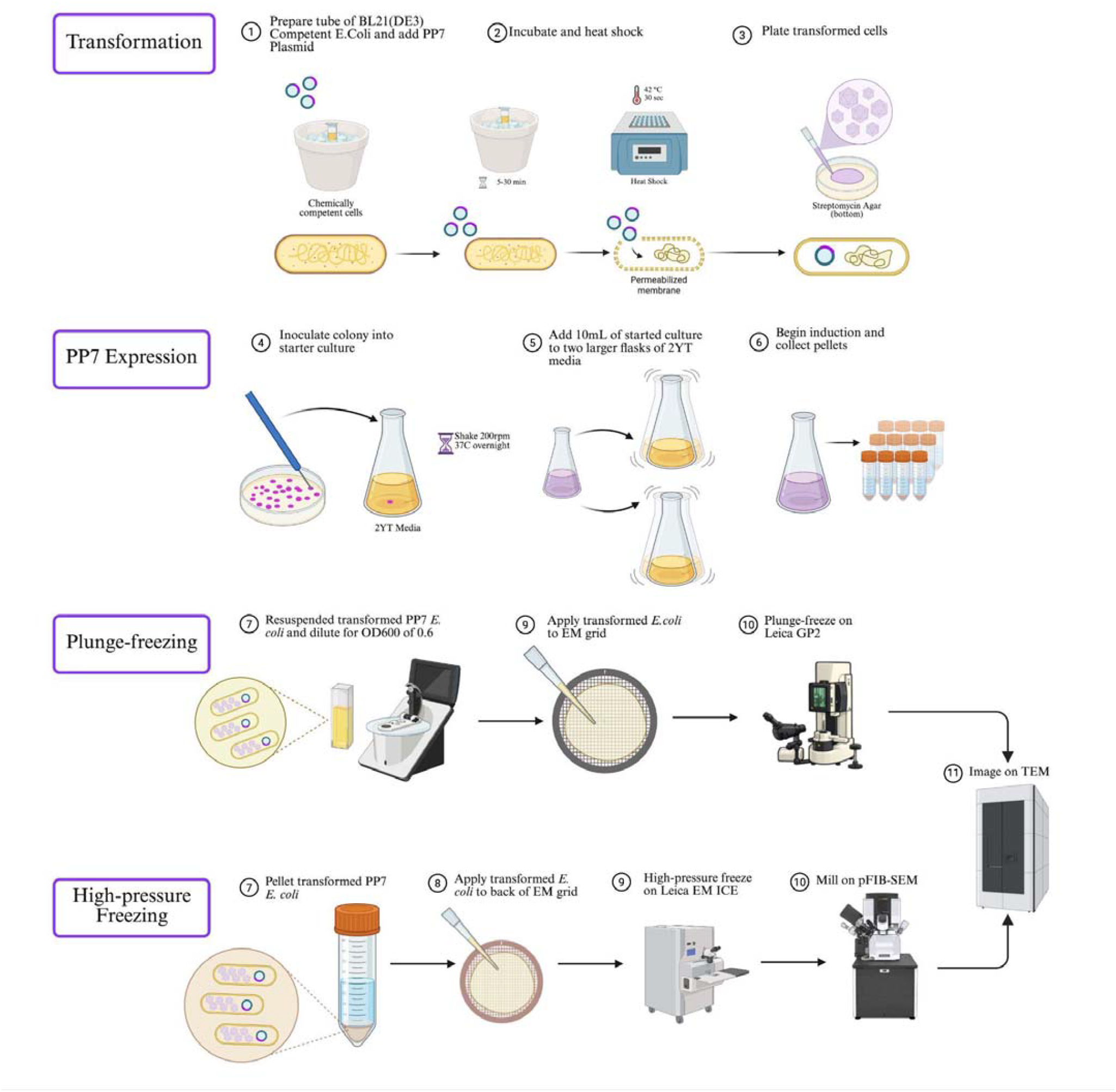
Expression of PP7 VLPs in *E. coli* and subsequent freezing. Four stages of preparing the VLPs in *E. coli* benchmark sample. With one round of transformation and expression, enough stock can be generated for tens of EM grids. These cells can either be plunge-frozen and directly imaged on the TEM without FIB-milling, or they can be high-pressure frozen and FIB-milled before imaging on the TEM.

**Extended Data Figure 6.**
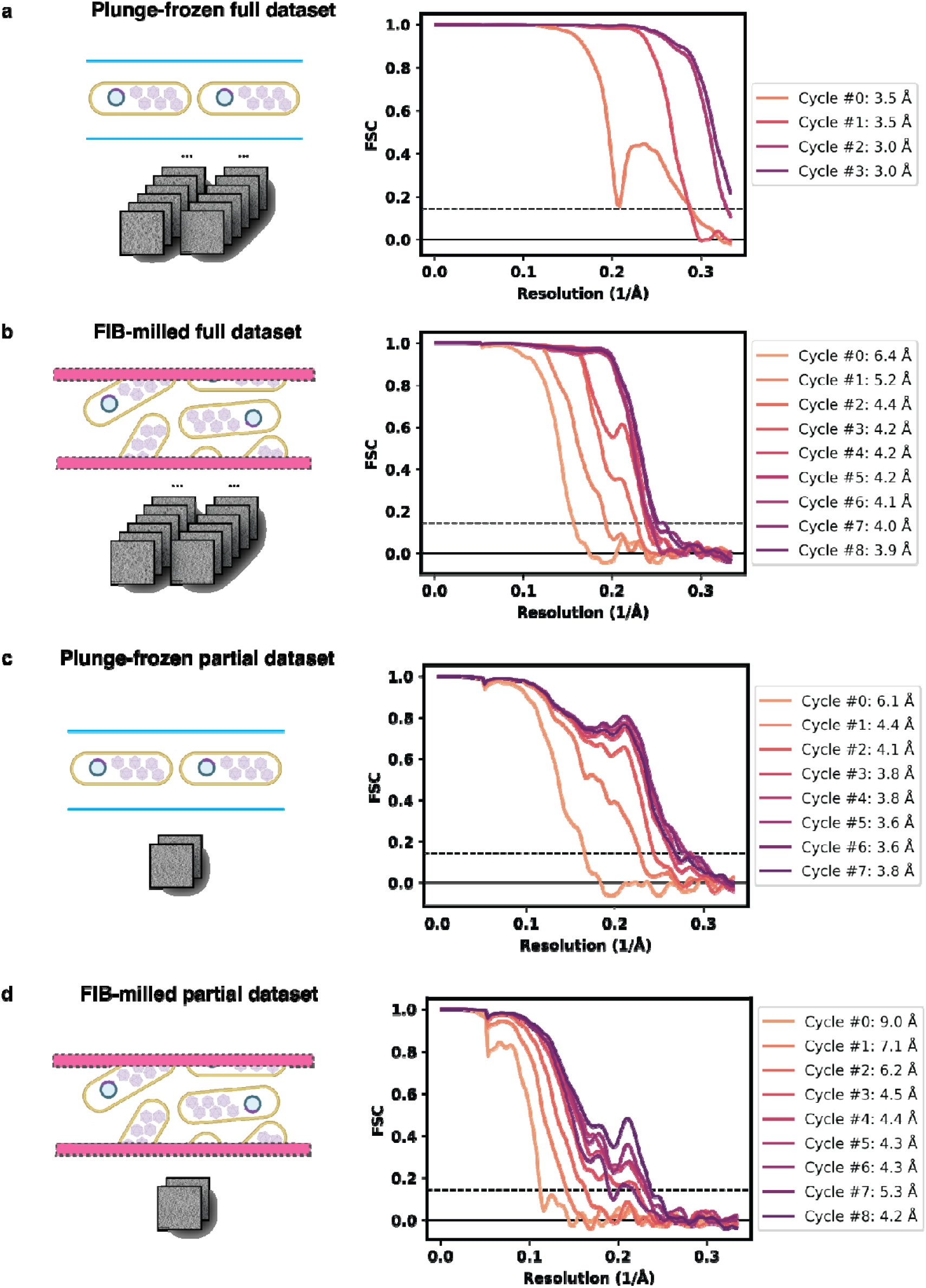
Stages of high-resolution STA refinements for full and partial datasets. FSC curves from stages of high-resolution refinements for full datasets of plunge-frozen (**a**) and FIB-milled (**b**) samples, and partial datasets for plunge-frozen (**c**) and FIB-milled (**d**) samples. One cycle refers to one iteration of CTF refinement, Bayesian polishing and 3D refinement. Cycle #0 is the last unbinned refinement prior to cycles of CTF refinement and polishing. All FSCs were calculated from the same soft-edge mask obtained from the VLP asymmetric subunit.

## References

1. Majumder, P. & Zhang, P. In situ cryo-electron microscopy and tomography of cellular and organismal samples. Curr. Opin. Struct. Biol. 93, 103076 (2025).

2. Tegunov, D., Xue, L., Dienemann, C., Cramer, P. & Mahamid, J. Multi-particle cryo-EM refinement with M visualizes ribosome-antibiotic complex at 3.5LÅ in cells. Nat. Methods 18, 186–193 (2021).

3. Russo, C. J., Dickerson, J. L. & Naydenova, K. Cryomicroscopy in situ: what is the smallest molecule that can be directly identified without labels in a cell? Faraday Discuss 240, 277–302 (2022).

4. Tuijtel, M. W. et al. Thinner is not always better: Optimizing cryo-lamellae for subtomogram averaging. Sci. Adv. 10, eadk6285 (2024).

5. Xing, H. et al. Translation dynamics in human cells visualized at high resolution reveal cancer drug action. Science 381, 70–75 (2023).

6. Peck, A. et al. AreTomoLive: Automated reconstruction of comprehensively-corrected and denoised cryo-electron tomograms in real-time and at high throughput. bioRxiv https://doi.org/10.1101/2025.03.11.642690 (2025) doi:10.1101/2025.03.11.642690.

7. Burt, A. et al. An image processing pipeline for electron cryo-tomography in RELION-5. bioRxiv https://doi.org/10.1101/2024.04.26.591129 (2024) doi:10.1101/2024.04.26.591129.

8. Khavnekar, S. et al. Optimizing Cryo-FIB Lamellas for sub-5Å in situ Structural Biology. bioRxiv https://doi.org/10.1101/2022.06.16.496417 (2022) doi:10.1101/2022.06.16.496417.

9. Lucas, B. A. & Grigorieff, N. Quantification of gallium cryo-FIB milling damage in biological lamellae. Proc. Natl. Acad. Sci. 120, e2301852120 (2023).

10. Berger, C. et al. Plasma FIB milling for the determination of structures in situ. Nat.Commun. 14, 629 (2023).

11. Keshavarz-Joud, P. et al. Pleomorphism in Wild-Type and Engineered PP7 Virus-Like Particles. Small 21, e06285 (2025).

12. Kelley, K. et al. Waffle Method: A general and flexible approach for improving throughput in FIB-milling. Nat. Commun. 13, 1857 (2022).

13. Klykov, O. et al. In situ cryo-FIB/SEM Specimen Preparation Using the Waffle Method. Bio-Protoc. 12, e4544 (2022).

14. Ermel, U. H. et al. copick: An open dataset interface and toolkit for collaborative annotation and analysis of cryo-electron tomography data. Protein Sci. 35, e70578 (2026).

15. Peck, A. et al. A realistic phantom dataset for benchmarking cryo-ET data annotation. Nat. Methods 22, 1819–1823 (2025).

16. Punjani, A., Rubinstein, J. L., Fleet, D. J. & Brubaker, M. A. cryoSPARC: algorithms for rapid unsupervised cryo-EM structure determination. Nat. Methods 14, 290–296 (2017).

16. Schwartz, J., Ermel, U.H., Ji, D., & Zhao, Z. Octopi: v.1.4. Zenodo (2026).

17. Schwartz, J., Ji, Di., Ermel, U.H., & Hutchings, J. py2rely: The Python Pipeline to Rely On. Zenodo. (2026).

